# Quantifying murine placental extracellular vesicles across gestation and in preterm birth data with tidyNano: a computational framework for analyzing and visualizing nanoparticle data in R

**DOI:** 10.1101/503292

**Authors:** Sean L. Nguyen, Jacob W. Greenberg, Hao Wang, Benjamin W. Collaer, Jianrong Wang, Margaret G. Petroff

## Abstract

Extracellular vesicles (EVs) are increasingly recognized as important mediators of intercellular communication that carry protein, lipids, and nucleic acids via the circulation to target cells whereupon they mediate physiological changes. In pregnancy, EVs are released in high quantities from the placenta and have been postulated to target multiple cell types, including those of the vascular and immune systems. However, most studies of pregnancy-associated EVs have used clinical samples and *in vitro* models; to date, few studies have taken advantage of murine models in which pregnancy can be precisely timed and manipulated. In this study, we used a murine model to determine whether the quantity of EVs is altered during healthy pregnancy and during inflammation-associated preterm birth. To facilitate data analysis, we developed a novel software package, tidyNano, an R package that provides functions to import, clean, and quickly summarize raw data generated by the nanoparticle tracking device, NanoSight (Malvern Panalytical). We also developed shinySIGHT, a Shiny web application that allows for interactive exploration and visualization of EV data. In mice, EV concentration in blood increased linearly across pregnancy, with significant rises at GD14.5 and 17.5 relative to EV concentrations in nonpregnant females. Additionally, lipopolysaccharide treatment resulted in a significant reduction in circulating EV concentrations relative to vehicle-treated controls at GD16.5 within 4 hours. Use of tidyNano facilitated rapid analysis of EV data; importantly, this package provides a straightforward framework by which diverse types of large datasets can be simply and efficiently analyzed, is freely available under the MIT license, and is hosted on GitHub (https://nguyens7.github.io/tidyNano/). Our data highlight the utility of the mouse as a model of EV biology in pregnancy, and suggest that placental dysfunction is associated with reduced circulating EVs.

## Introduction

Extracellular vesicles (EVs) encompass a broad class of membrane-enclosed structures secreted by cells, and are classified based on their size and subcellular origin. Exosomes, which range from 40-150nm, arise from the inward budding of late endosomes, and microvesicles, which range from 100-1000nm, develop as a result of outward budding of the plasma membrane. Because EVs have the capacity to induce physiological responses in recipient/target cells, they are of immense interest for many life science disciplines including immunology, cancer biology, and medicine [1–4]. Molecular contents of EVs, which include lipids, proteins, and nucleic acids, serve as the basis for intercellular communication [5]. Indeed, EVs have been shown to be effective and important in mediating processes including antigen cross-presentation [6], establishing local ‘niches’ for metastasis of cancer cells [7], delivery of microRNAs for suppression of gene expression in target tissues [8], and even transfer and subsequent translation of mRNAs into target cells [9].

Within the context of pregnancy, the conceptus-derived placenta secretes copious amounts of EVs that are detectable in maternal blood as early as the first trimester in women [10–12]. Humans and mice share in common the hemochorial anatomic arrangement of the placenta, in which maternal and fetal circulations are separated by only 2 to 4 cellular layers. Importantly, in both species, the embryo-derived trophoblast is suffused with maternal blood during much of pregnancy, serving as a surrogate ‘endothelium’ across which maternal nutrients and fetal waste products are exchanged. This intimacy also allows for the trophoblast to shed large amounts of EV directly into the maternal circulation. Studies in women have suggested that placental EVs contribute to critical processes in pregnancy, including vascular development of the maternal-fetal interface and establishment of maternal immune tolerance to the antigenically foreign fetus [1,13]. Further, because placental EVs circulate in maternal peripheral blood, they can serve to provide a noninvasive ‘liquid biopsy’ indicating fetal health in utero; similar concepts are applied to overall health as a potential diagnostic for cancer [3,7]. The similarities in placentation between humans and mice highlight the utility of the murine model to understand the function of pregnancy-associated EVs; surprisingly, however, only a few recent studies [14,15] have explored the pregnant mouse as a model for placental EV function, and no studies have yet explored the response of pregnancy-associated EV to inflammation-induced preterm birth.

Interest in EVs has burgeoned for the above reasons; however, their isolation and analysis has posed a number of technical challenges. Because of their small size, direct observation and quantification of isolated EVs require specialized equipment that generates large amounts of data. One method of measuring the size and concentration of nanoparticles is nanoparticle tracking analysis (NTA; NanoSight, Malvern Instruments, USA), which makes use of a microfluidic system and laser together with a camera and software to track individual particles, measuring their size and concentration on the basis of Brownian motion [16,17]. However, while NanoSight provides an effective means of measuring EVs, experiments with more than one condition require the use of independent spreadsheet software for extraction of raw counts, treatment information, statistical analysis, and graphical representation. Further, because this approach necessitates direct manipulation of raw data by the user, it is susceptible to user- or software-introduced error [18]. Analysis of large number of samples would benefit from a computational approach but requires experiment-specific scripts, which are not easily adaptable to complex experimental designs or conducive for reproducible research [19, 20].

In this study, we aimed to establish baseline quantities of murine EVs across pregnancy, and further to determine the effects of inflammation on pregnancy-associated EVs in a model of preterm birth. In addition, we sought to develop a computational framework to standardize the process of NTA analysis including data import, organization, visualization, and statistical analysis.

## Materials and methods

### Mice and treatments

All experiments were approved by Michigan State University Institutional Animal Care and Use Committee protocol: 04/18-050-01. Mice (C57Bl/6; Jackson Laboratory, Bar Harbor, ME, USA) were anesthetized by 3% isoflurane and euthanized by cervical dislocation and subsequent bilateral pneumothorax. Mice were housed in temperature-controlled environments in a 12-hour light/dark cycle with standard diet and water available ad libitum. Timed matings were performed in 6-8 week old mating pairs, and the presence of a vaginal plug was designated as gestational day (GD) 0.5. For preterm birth experiments, mated GD16.5 females were injected with 10mg LPS (Salmonella enterica, Sigma-Aldrich, St. Louis, MO, USA) in 100μl PBS or 100μl PBS (vehicle control) i.p. and sacrificed 4 h later. Following anesthesia with isofluane and oxygen (2L/min), whole blood was harvested via cardiac puncture using 1.2ml S-Monovette EDTA-containing tubes fitted with a 22-gauge needle (Sarstedt, Newton, NC, USA). Plasma was isolated by two rounds of centrifugation at 2000 x g for 15 minutes at 4°C and frozen at −80°C.

### Exosome isolation and validation

Frozen plasma samples were thawed on ice and exosomes were harvested using Total Exosome Isolation reagent (Thermo Fisher, Burlingame, CA, USA) for plasma following the manufacturer’s instructions. Samples were resuspended in 25-50 μl of phosphate buffered saline (PBS) [Corning, Manassas, VA, USA]. Plasma exosomes were examined by transmission electron microscopy as previously described [2] (S1 Fig). Briefly, pelleted exosomes were resuspended with 4% paraformaldehyde in PBS, loaded on formvar-carbon coated grids and counterstained with 2.5% glutaraldehyde and 0.1% uranyl acetate in PBS, and imaged on a JEOL100 CXII (JEOL, Peabody, MA, USA).

### NanoSight analysis

Nanoparticle tracking analysis was performed on a NanoSight NS300 (Malvern Panalytical, Westborough, MA, USA) equipped with a 488nm excitation laser and an automated syringe sampler. Plasma EV samples were diluted 1:125-1:500 in PBS and loaded into 1ml syringes with the flow rate set to 50 and the operator blinded to sample identity. Diluted samples were measured by two separate injections, each by three 30-second videos. Measurements were recorded at camera level 11 and detection threshold of 4. Experiment summary CSV files generated by NTA software v3.2 and used for computational analysis and development of software.

### tidyNano Software development

Fig 1 summarizes the core functions of tidyNano, which serve to simplify importation of raw NanoSight data into a data frame suitable for rapid computational analysis in the R environment. To this end, tidyNano rearranges data into a ‘tidy’ format in which observations are represented in rows and variables in a single column, rearranging the data from the canonical ‘wide’ format (provided by NTA software), to the ‘long’ format that is characteristic of ‘tidy’ data (Fig 2, 3A, 3B) [21], and conducive for manipulation with the dplyr package [22], visualization with the ggpplot2 package [23], and computation with base R statistics [24] (Fig 1). tidyNano also parses out sample information and performs back calculations to account for sample dilution during NTA measurement. Once in this ‘tidy’ format, tidyNano provides functions to summarize data and calculate statistics that are immediately suitable for further analysis visualization with ggplot2 and/or shinySIGHT, as described below (Fig 1). Functions within tidyNano make use of dplyr and tidyr packages to transform and aggregate data, allowing each tidyNano output to be compatible with a wide variety of R packages and suitable for recent changes in visualization paradigms [22,23,25,26].

**Fig 1.**
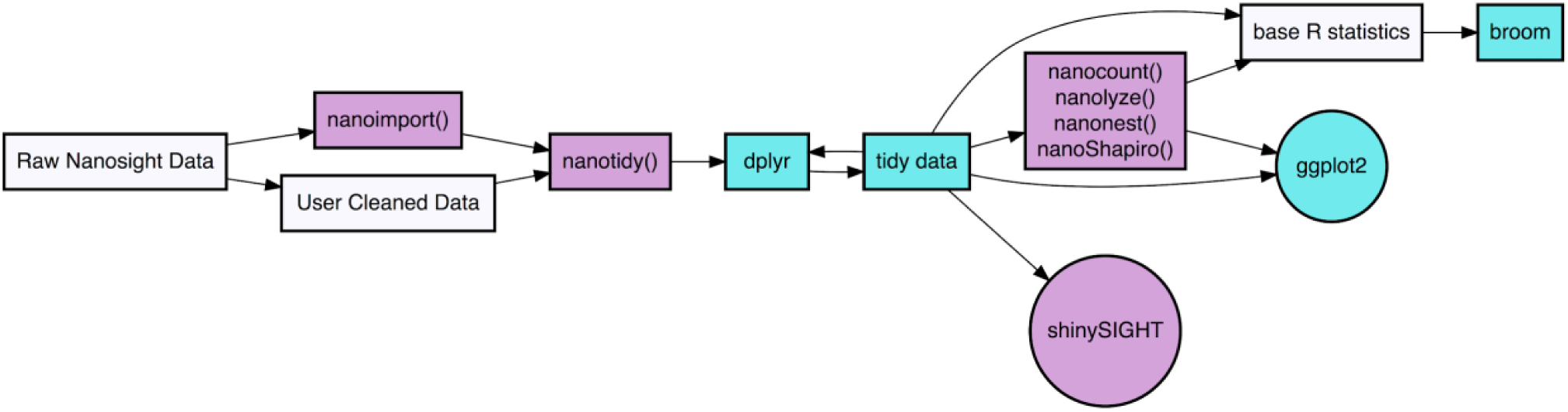
Schema of tidyNano framework. tidyNano (purple) is designed to facilitate the process of importing and formatting data into a tidy format, such that the data are compatible with existing visualization and statistical packages (cyan).

**Fig2.**
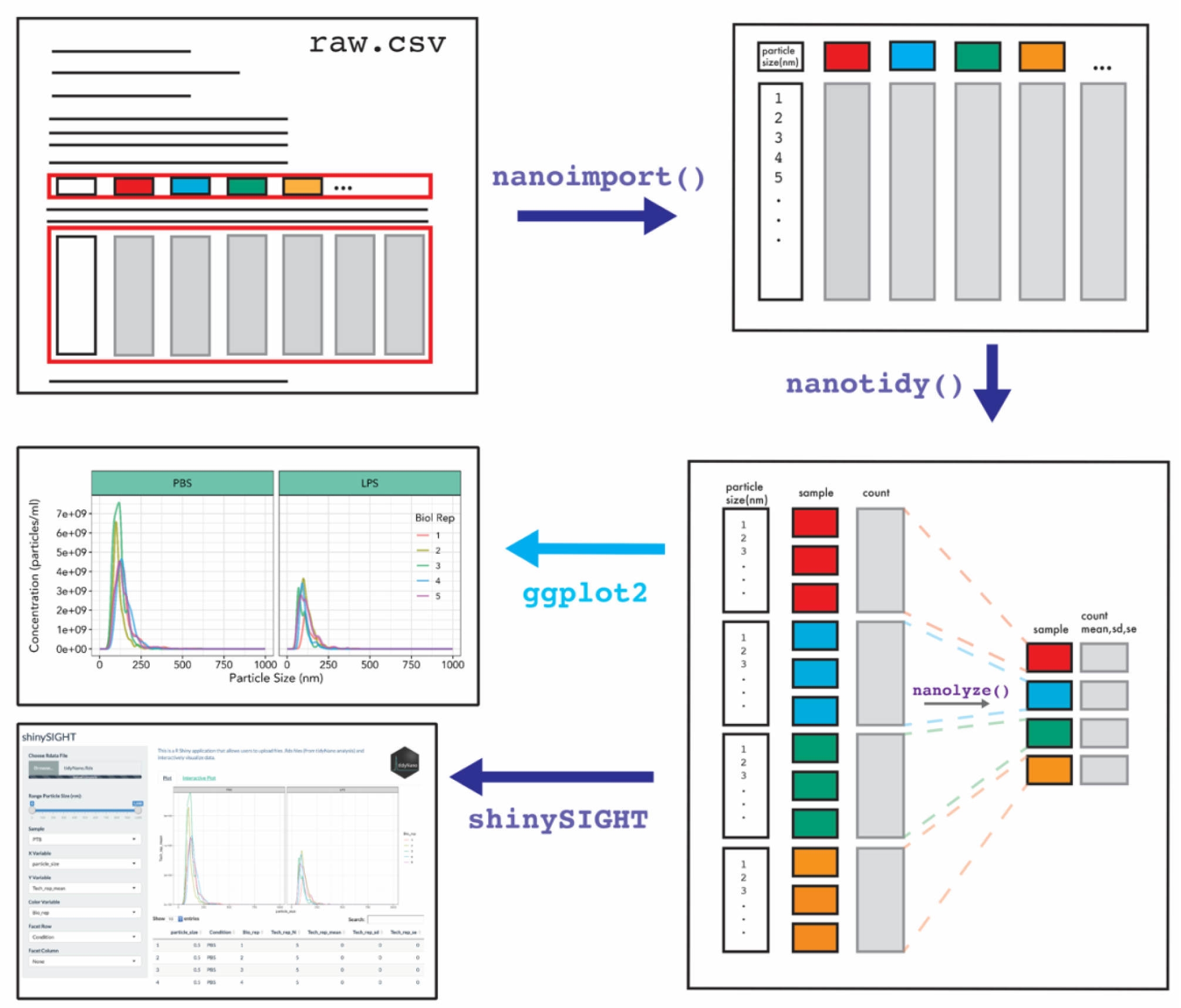
Example workflow of tidyNano for analysis of NTA data. Core functions of tidyNano (violet) are to facilitate extraction, formatting and aggregation of NTA data. Following data import, tidyNano functions can be easily visualized using existing packages such as ggplot2 or with the interactive web application shinySIGHT.

**Fig 3.**
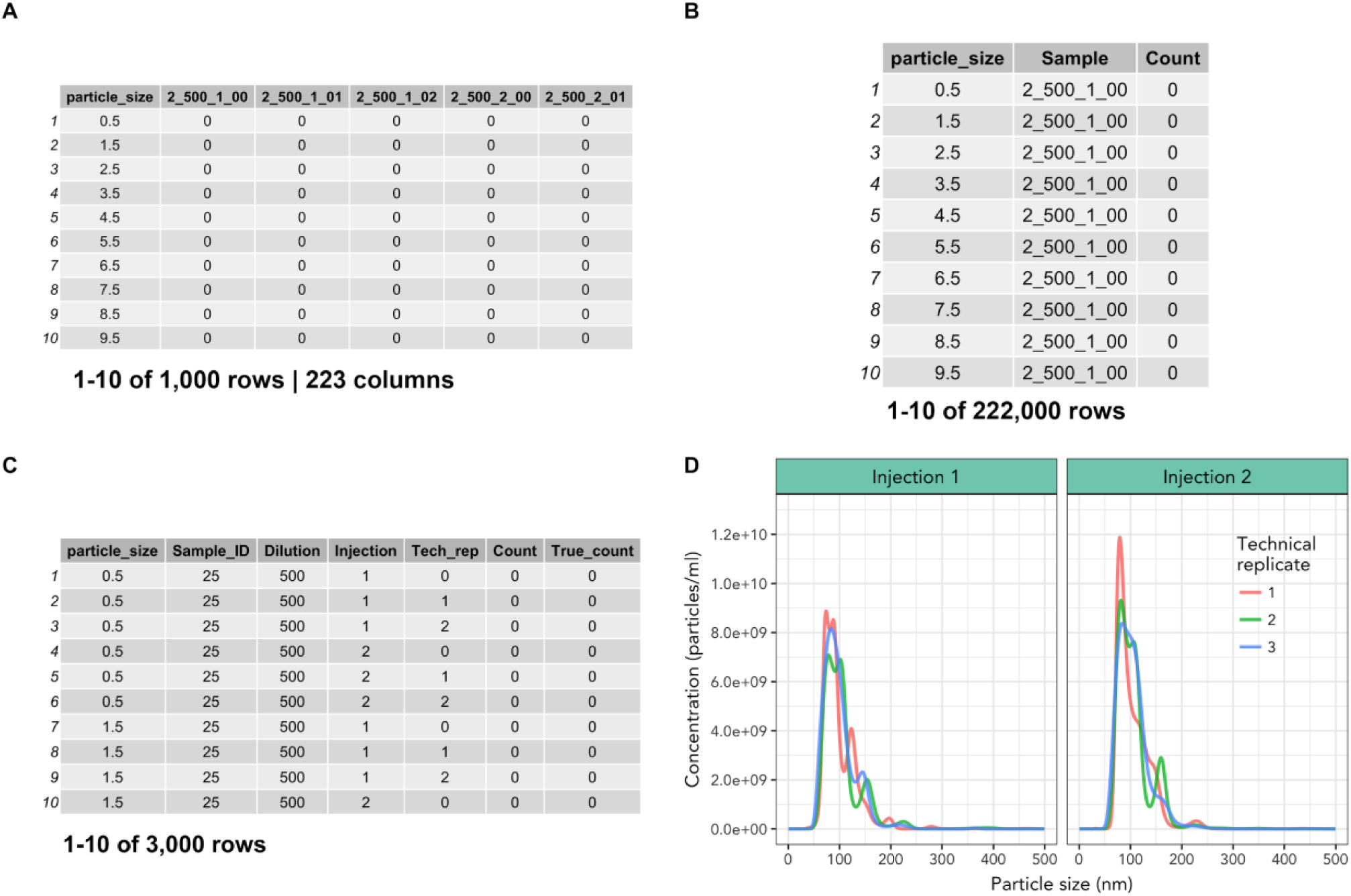
Data import and reformatting with nanoimport() and nanotidy(). (A) Output from nanoimport() or manually-cleaned NTA data. (B) Intermediate step of nanotidy() function which converts data from ‘wide’ to ‘long’ format, generating tidy data that can be easily filtered to isolate individual sample values. (C) Finalized output from nanotidy() function which separates the ‘Sample’ column into user-specified columns. (D) Representative visualization of technical replicates of nanoimport() output which includes multiple sample injections from one sample with ggplot2, lines represent technical replicate measurements.

### Data import and cleaning

TidyNano provides nanoimport() and nanocombine() functions to import individual NTA experiment files or combine multiple experiment files into a single data frame within the R environment. Both functions determine NTA software versions and import raw tabular data, and can accommodate user-specified NTA parameters including bin widths and particle ranges. The functions automatically determine the number of samples in an NTA experiment, extract sample names and particle counts for each sample, and return a single data frame. To do this, the nanotidy() function uses the gather() function from the tidyr package to convert the data from ‘wide’ to ‘long’ format, i.e., the condensing of individual sample columns into key-value pairs [21]. The function then splits the sample column into multiple columns based on user-specified names, and performs back calculations to determine the true count of particles in the original sample. Finally, the column headers are converted to the correct class (e.g., categorical factor or numeric) (Fig. 2).

### Data summarization

The nanolyze() function calculates summary statistics by group and generates a data frame that includes the number of samples within groups, mean, standard deviation, and standard error. Further, it calculates a wide variety of statistics such as technical and biological replicate means or differences between other experimental conditions. Nanolyze() contains arguments for specifying prefixes to summary statistic columns which allows for the function to be used sequentially to average replicates such as technical, biological, and group/treatment replicates. Each iteration of Nanolyze returns a visualizable tidy data frame (Fig 2). The nanocount() function allows for rapid calculation of the total sum of particles within groups of samples and can be combined with existing functions such as filter() to subset data.

### Data visualization and statistical analysis

We also developed shinySIGHT, a web application built within the R shiny framework that allows the user to upload, interact with, and visualize the NanoSight data without needing to program. shinySIGHT facilitates dynamic exploration of NTA data and can provide an understanding of particle size distribution and concentration with user-adjustable sliders to specify particle size ranges (Fig 2). shinySIGHT is available within the tidyNano package and can be run locally by using the shinySIGHT() command.

## Results

To demonstrate the utility of the package, we used tidyNano to analyze three NanoSight datasets. The first set consisted NanoSight data from polystyrene fluorescent and non-fluorescent bead standards (S2 Fig) and a second set consisted of peripheral exosomes from C57Bl/6 female mice across six time points of pregnancy. The third dataset consisted of peripheral exosome data from a lipopolysaccharide (LPS) model of preterm birth in GD16.5 C57 Bl/6 mice.

### Data Import and Cleaning

To demonstrate the nanotidy() function, we imported raw NTA data from a dataset of murine plasma exosomes from a preterm birth model of pregnancy. We used nanoimport() to import NTA data into R which worked by extracting column headers and individual particle counts and combined them into a large data frame (S3 Fig). If desired, NTA data can also be cleaned manually by the user in spreadsheet software such as Excel, as was done with the murine pregnancy time course exosome dataset (Fig 3A). Next, nanotidy() was used to reshape the data into a key-value pairs such that the individual samples and values were condensed into single columns resulting in the conversion from ‘wide’ to ‘long’ format in an intermediate step (Fig 3B). Nanotidy() then parsed the ‘Sample’ column into individual (user-specified) columns and performed back-calculations to obtain a ‘True_count’ column containing the corrected count values (Fig 3C). The output of nanotidy() resulted in a tidy data frame that could be summarized, visualized or manipulated for further analysis (Figs 1 and 2).

### Data summarization

To study the concentration of peripheral exosomes in murine pregnancy, we sampled blood from 6 experimental groups (nonpregnant, four time points of gestation corresponding to different embryonic developmental stages, and one postpartum) (n=5-7/group). From each experimental group, three technical replicates, and two injection replicates were measured, resulting in 222 distinct measurements (S4 Fig). Nanolyze() was used to average the technical and injection replicates within each biological replicate, resulting in a data frame that was suitable for plotting (S5 – S6 Fig). To demonstrate sequential use of nanolyze(), we calculated averages of technical replicates from two separate injections of the same dilution (S5 Fig, Fig 4A). Nanolyze() was then repeated sequentially to determine the mean exosome concentration between biological replicates using data in Fig 4A as input (Fig 4B). This was possible due to the name argument, which allows custom naming of the column headers by users, and param_var argument, which specifies the column on which to perform the calculation. We then used nanocount() (data in Fig 4A) to determine the total concentration of particles within each biological replicate, resulting in a data frame that could be plotted.

**Fig 4.**
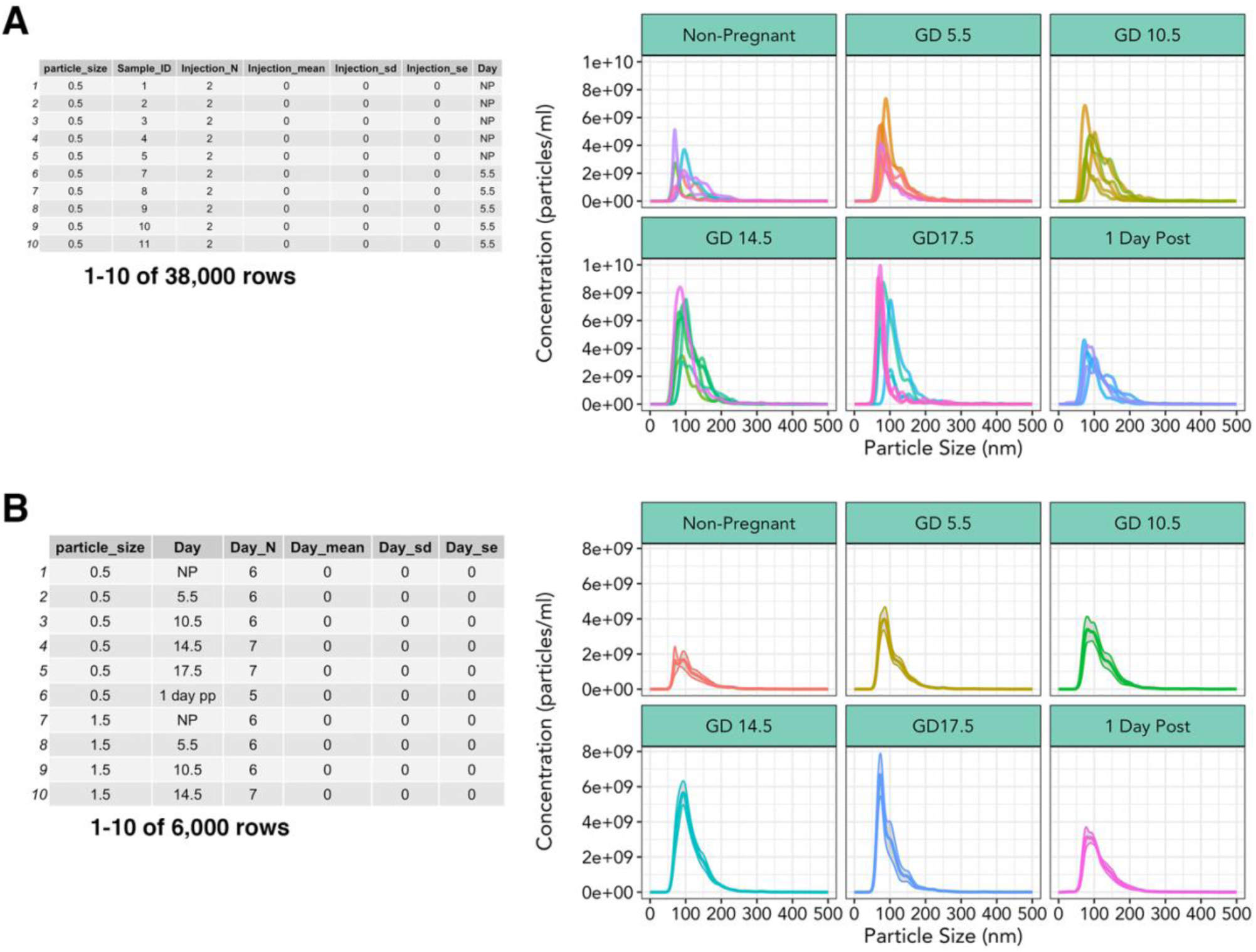
Multiparameter summary statistics and visualization. (A) Output from nanolyze() which calculates mean, standard deviation, and standard error within specified groups. Corresponding line graph of exosome concentration as a function of size, split by gestational age, different colored lines within each group represent a single biological replicate. (B) Summarization of biological replicate data from nanolyze() of 3(A) data. Corresponding line graph depicting mean exosome concentration of biological replicates as a function of size, grey area surrounding lines represent standard error of the mean.

### Visualization

Each tidyNano function described above returns a data frame conducive for flexible downstream graphical representation. In the examples above, we created all plots from nanolyzed data using ggplot2. Further, we developed an R Shiny web application, shinySIGHT, that facilitates NTA data visualization, interactive exploration, and interpretation, and may be particularly practical for users who are less experienced in computer programming. To this end, after importing and tidying the murine exosome time course data with nanoimport() and nanotidy(), respectively, we used the nanosave() function to create a .Rds file. We then used the shinySIGHT() function to launch shinySIGHT, and imported the .Rds file. This allowed interactive visualization of NTA data from the preterm birth data without needing to be explicitly coded (Fig 5). A detailed vignette outlining how to use shinySIGHT is available at https://nguyens7.github.io/tidyNano/articles/tidyNano.html#tidynano-vignette.

**Fig 5.**
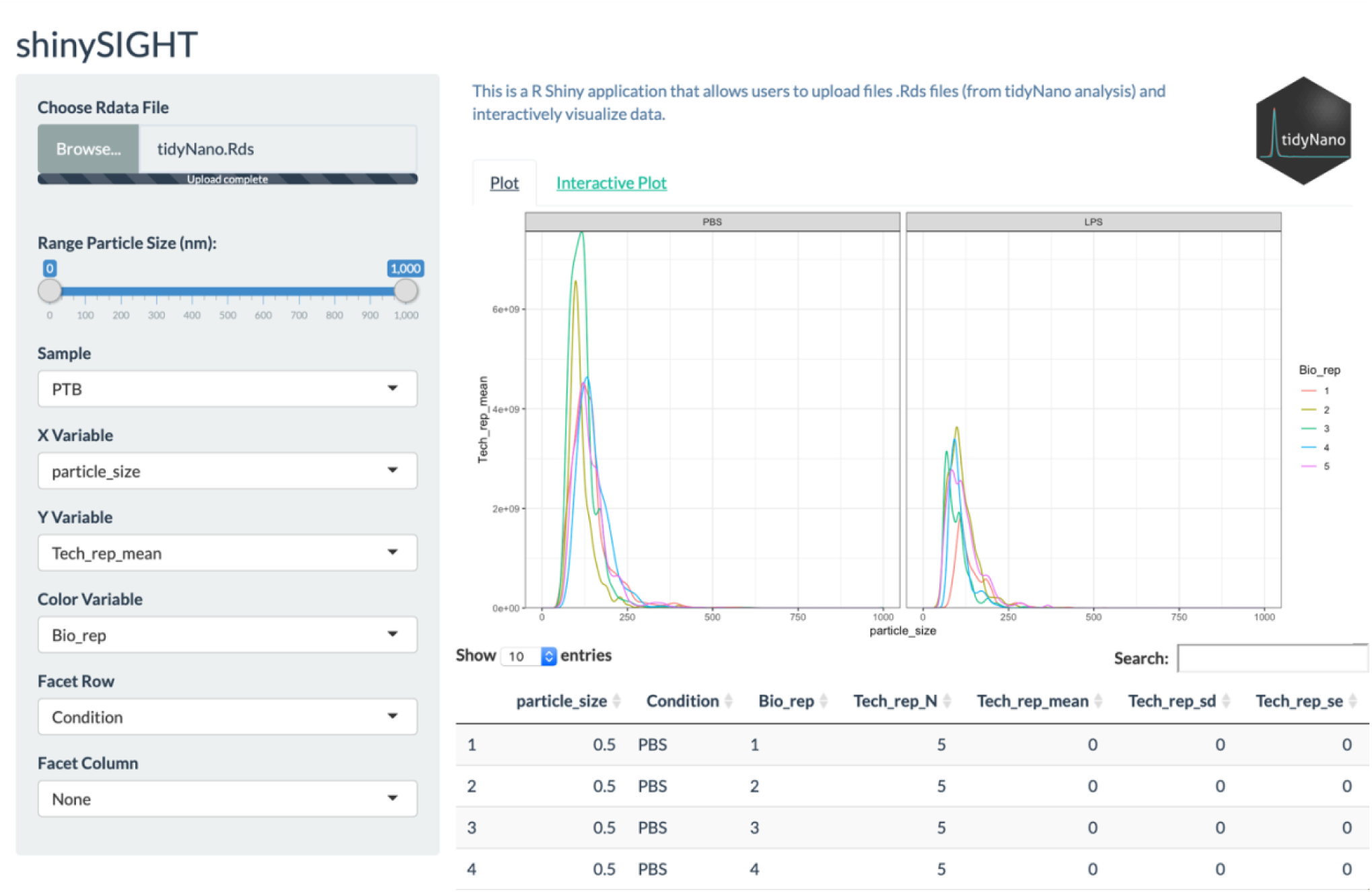
Interactive data manipulation and visualization with shinySIGHT web application. shinySIGHT allows users to upload tidyNano data to visualize and manipulate data using a graphical user interface. shinySIGHT automatically generates plots from user uploaded data as well as displays the underlying data frames that make up the visualizations.

If desired, technical and injection replicates (S5 Fig) and/or individual samples (Fig 3D) may be visualized using shinySIGHT (not shown). Summarization steps from each time point across gestation was examined by line graph using ggplot2, and demonstrated mean exosome concentration of each animal split by gestational time point (Fig 4A) as well as mean concentration of biological replicates (Fig 4B). After plotting mean concentration of individual replicates with nanocount(), total exosome concentration was plotted to determine the variations in exosome concentrations across gestation (Fig 6B).

**Fig 6.**
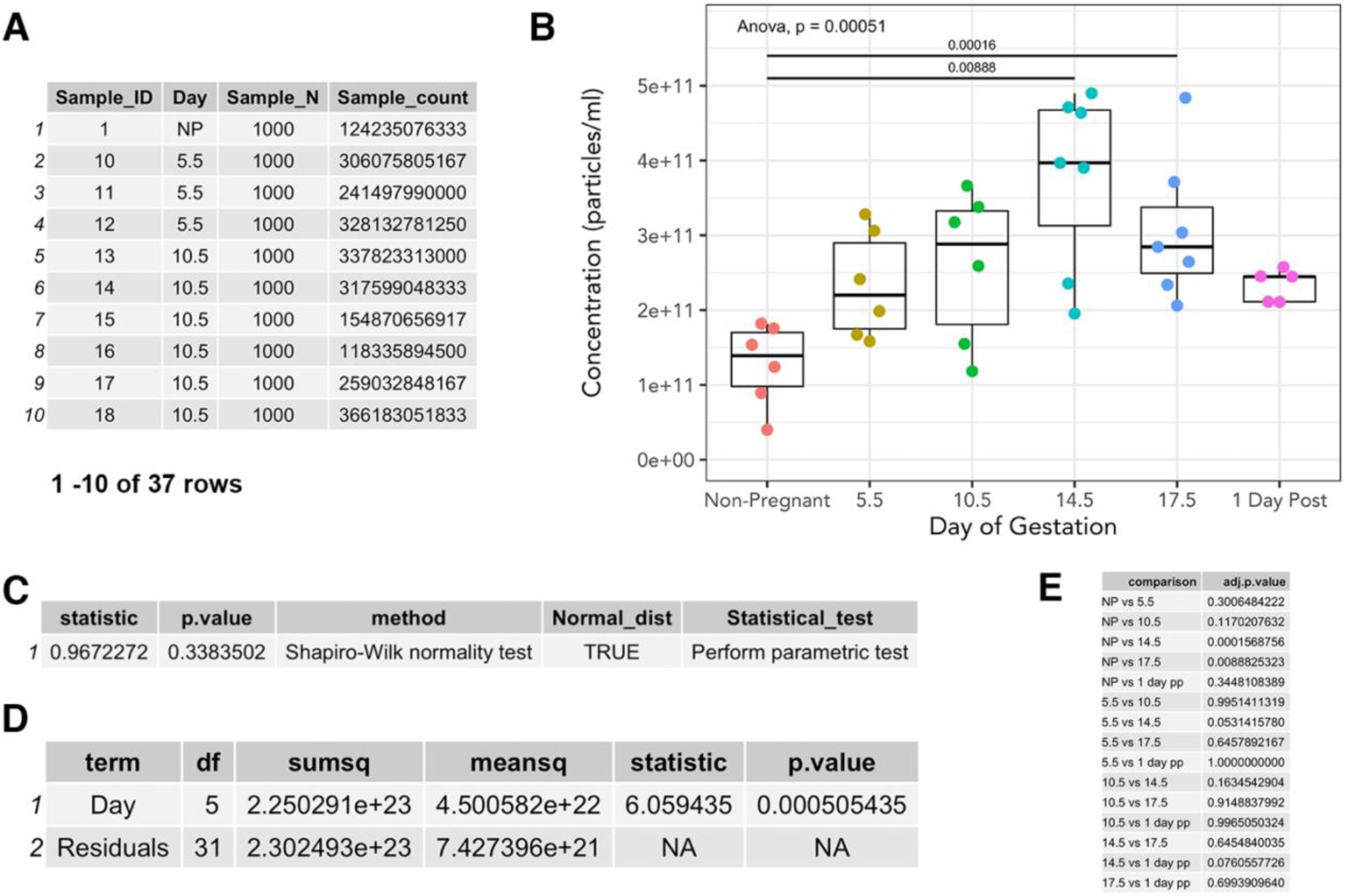
Calculation of extracellular vesicle counts and statistics with nanocount(), nanoShapiro() and nanoTukey(). (A) nanocount() function determines the total concentration of particles. (B) Boxplot of murine peripheral exosomes across pregnancy, each point represents an individual biological replicate. (C) nanoShapiro() function determines Gaussian distribution of data. (D) ANOVA output generated from nanocount of data frame in (A). (E) nanoTukey() output of pair-wise Tukey post hoc analysis comparing extracellular vesicle concentration across gestation.

### Statistical analysis

We used nanoShapiro() to determine the normality of the murine exosome dataset, which returns a data frame with suggestions for the appropriate statistical test (Fig 6C). Next, we used existing base R statistical packages [24] to run an ANOVA on our samples to compare total exosome count across gestational days using data in Fig 6A. The result of the ANOVA suggested that there were significant differences in exosome concentrations across gestational age (p < 0.001) (Fig 6D). We then utilized the nanoTukey() function to run a Tukey post-hoc test, which returned a tidy data frame of pair-wise comparisons of exosome concentrations (Fig 6E).

### Effects of gestation day and inflammation on circulating EV concentrations

Development of the nanoTidy package enabled rapid analysis of the changes in circulating EVs across gestation (Fig 6B) and following LPS treatment (Fig 7). Maternal plasma EV rose linearly with advancing gestational age, peaking at GD14.5 and corresponding with placental mass (S7 Fig). EV concentrations were significantly higher (p < 0.05) between non-pregnant and GD14.5, non-pregnant and GD17.5, and trends (p < 0.1) towards an increase between GD5.5 and GD14.5. Likewise, there was a trend towards reduction in circulating EVs between GD14.5 and 1 day postpartum. Further, EV concentrations at GD5.5 and 10.5 were intermediate between those of nonpregnancy and GD14.5 and 17.5.

**Fig 7.**
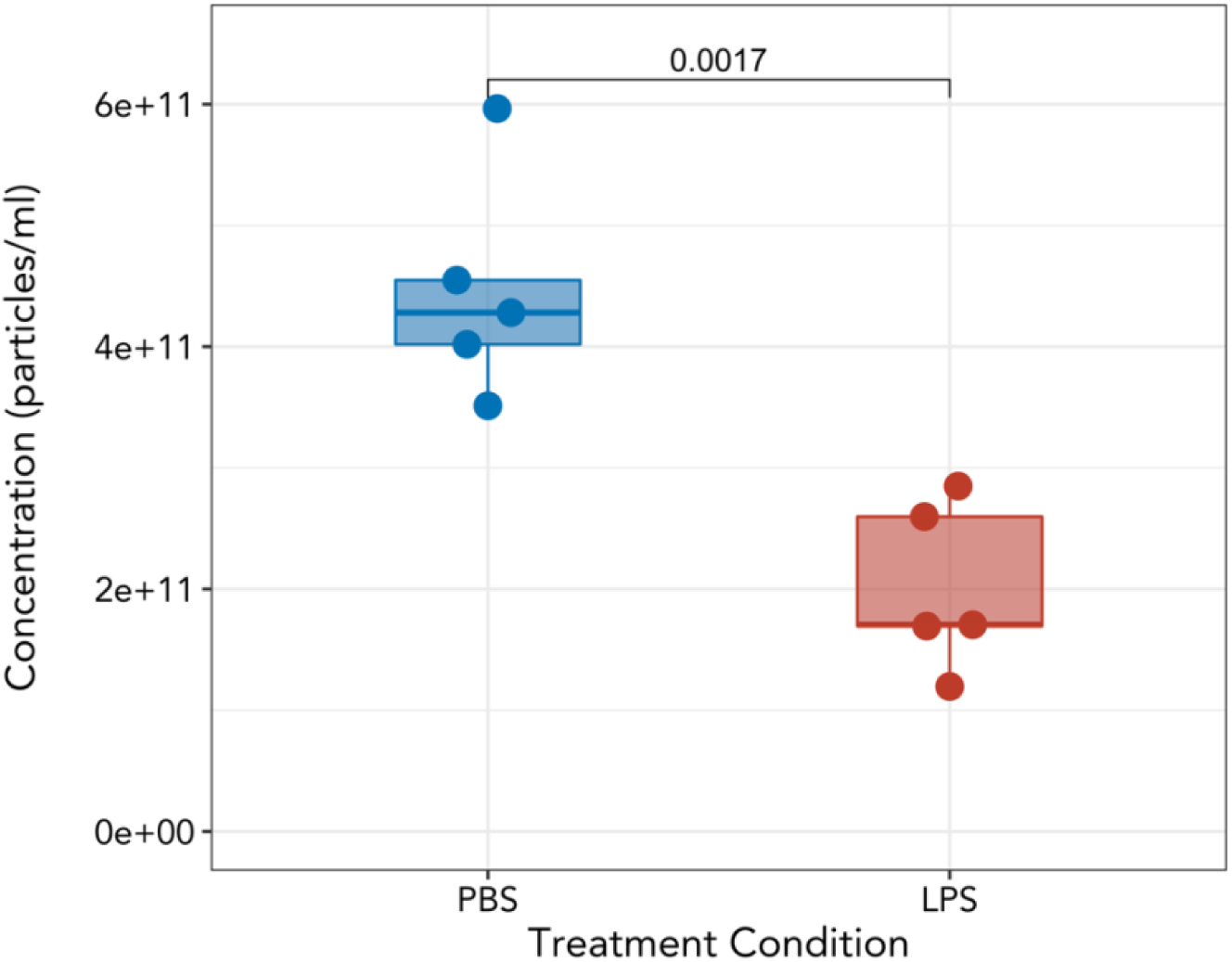
Peripheral exosome concentration of GD16.5 mice treated with 10ug LPS. Plasma samples were collected 4 h following i.p. injection of phosphate buffered saline (PBS) or 10μg LPS. Each point represents a biological replicate. Welch’s t-test.

Administration of 10μg LPS i.p. in GD16.5 mice resulted in wholesale fetal death as indicated by hemorrhagic implantation sites, as early as 4 hours post-treatment and indicative of impending preterm delivery. This was associated with a rapid and significant decrease circulating EV concentration (p < 0.0017) (Fig 7).

## Discussion

In this study, we quantified the murine plasma EVs across normal gestation, as well as in a model of inflammation-induced preterm birth. The principle findings were that plasma EVs concentrations increased with advancing gestational age in mice in comparison to nonpregnant females, and that LPS-induced fetal loss was associated with a striking reduction in circulating EV. Additionally, we report a novel pipeline in the R platform for rapid exploration and visualization of data generated from NanoSight analysis of EVs.

In pregnant females, plasma EV concentrations rose progressively and were highest during the latter stages of gestation, reaching a maximum at GD14-17. This finding is consistent with both a recent report of EV across gestation in mice and with reports that EVs are highest during the third trimester of human pregnancy [14,27]. Additionally, mean plasma EV concentrations at GD5.5 and 10.5 rose to intermediate concentrations levels between those of nonpregnancy and GD14.5, reminiscent of increases in circulating EV observed during the first trimester of human pregnancy [27]. This may reflect heightened release of maternal, rather than placental, EVs, as neither the first trimester of human pregnancy nor the first half of murine pregnancy is characterized by established maternal blood flow to the placenta. Factors regulating an early pregnancy-associated increase in EV are unknown; hormones from the conceptus and embryo are already very high at these stages, and may induce systemic – possibly vascular – release of maternal EVs. Early rises in circulating EV during pregnancy may reflect a mechanism that controls systemic changes in the mother in preparation for her own dramatic physiological adaptations to pregnancy and/or her accommodation of rapid fetal growth.

EV concentrations tended (p = 0.076) to drop rapidly at one day postpartum from the highest values at GD14. These data are also in line with rapid clearance of placental EVs from the circulation of women following birth [10]. Changes in exosome content during late gestation have been suggested to contribute to parturition in mice, due in particular to a progressive increase in EVs carrying proinflammatory mediators [14]. Alterations in the EV microRNA profile towards late gestation were also observed in women, and further changes occurred in preterm birth [28]. Interestingly, we observed a striking reduction in circulating EV concentrations following LPS administration into pregnant dams, modeling infection-induced preterm birth. This reduction in peripheral exosomes may reflect acute placental dysfunction and thus placental EV secretion. In humans, peripheral EVs increase in pre-eclampsia, however no quantitative differences were observed between normal term and spontaneous preterm birth [28–30]. Time of sample collection relative to onset of preterm birth symptoms as well as variable etiology of preterm birth in human clinical samples may explain differences observed in our study. It also remains possible, and even likely, that inflammation-induced preterm birth in mice alters EV cargo as in women. Ongoing studies are addressing this possibility.

In addition to our experimental findings, we developed a computational framework, tidyNano, in the R platform that facilitates analysis of data generated by NanoSight. Following sample measurement, the NanoSight software generates a PDF report that includes a line graph depicting size distribution and particle concentration, as well as summary statistics including mean, median, and mode size of the EVs (S8 Fig). For downstream analysis, including combinatorial analysis of multiple datasets within a single experiment, users can access the raw NanoSight data, which includes particle counts and size, as well as user-defined acquisition parameters. However, experiments typically consist of multiple conditions, and require the use of independent spreadsheet software (e.g., Microsoft Excel). Although spreadsheets can facilitate this by providing intuitive processing through a point-and-click interface, the requirement for users to directly interact with raw data is a major disadvantage as it is tedious and increases the chances for user-introduced error [18]. Further, Excel itself has been known to inadvertently change underlying raw data (e.g., the gene name OCT4 can be converted to the date 10-04), which can also become problematic for downstream analysis [18,31,32]. Spreadsheet software is also limited to user-specific pipelines, cannot be easily expanded to accommodate multiparameter experiments, and the work and time required to analyze data typically increases linearly with added data.

To develop software that avoids these pitfalls, we took advantage of a recent paradigm in computational data analysis known as data ‘tidying’, which has become increasingly popular due to its consistent format that is amenable to rapid data exploration and analysis [21]. Tidy data is organized into a format such that observations are represented in rows and variables in a single column, which allows for subsetting, statistical calculation, and visualization of the data. Several popular packages that accept tidy data have been developed, and allow for efficient rearrangement, manipulation, programming, and visualization of data [22,23,25,26]. TidyNano employs a similar strategy, allowing users to quickly import and transform nanoparticle data into a ‘tidy’ format. We showed here that this software was able to efficiently process and analyze large sets of biological data. We also created an R Shiny web application, shinySIGHT, for further exploration of the data using a graphical user interface that can facilitate the process of inspecting and visualizing data without requiring the user to know how to code.

In summary, the present study reveals changes in EV concentrations across gestation and in a model of inflammation-induced preterm birth, and describes a novel software package that provides a framework for analyzing NTA data by performing functions that address the key components of data analysis: import and cleaning, summarization, visualization, and statistical calculation. TidyNano, which will be updated to accommodate future NanoSight models and NTA software, is an open source package developed in R, hosted on GitHub (https://github.com/nguyens7/tidyNano), and is freely available under the MIT license. Data, supporting documentation, and vignettes can be accessed at the package website (https://nguyens7.github.io/tidyNano/). TidyNano summarization functions are general purpose and can be adapted to analyze other tidy data, including non-nanoparticle data.

## Acknowledgements

The authors thank Alicia Withrow for transmission electron microscopy analysis assistance, and Gregory Burns and Thomas Spencer of the University of Missouri, Wei-Ting Hung and Lane Christenson of The University of Kansas Medical Center, and Neva Kandzija and Manu Vatish of The University of Oxford for validation and feedback on the tidyNano package. The authors also acknowledge Soo Hyun Ahn and Sarika Kshirsagar for helpful discussions in designing and implementing the software.

---

**S1 Fig.**
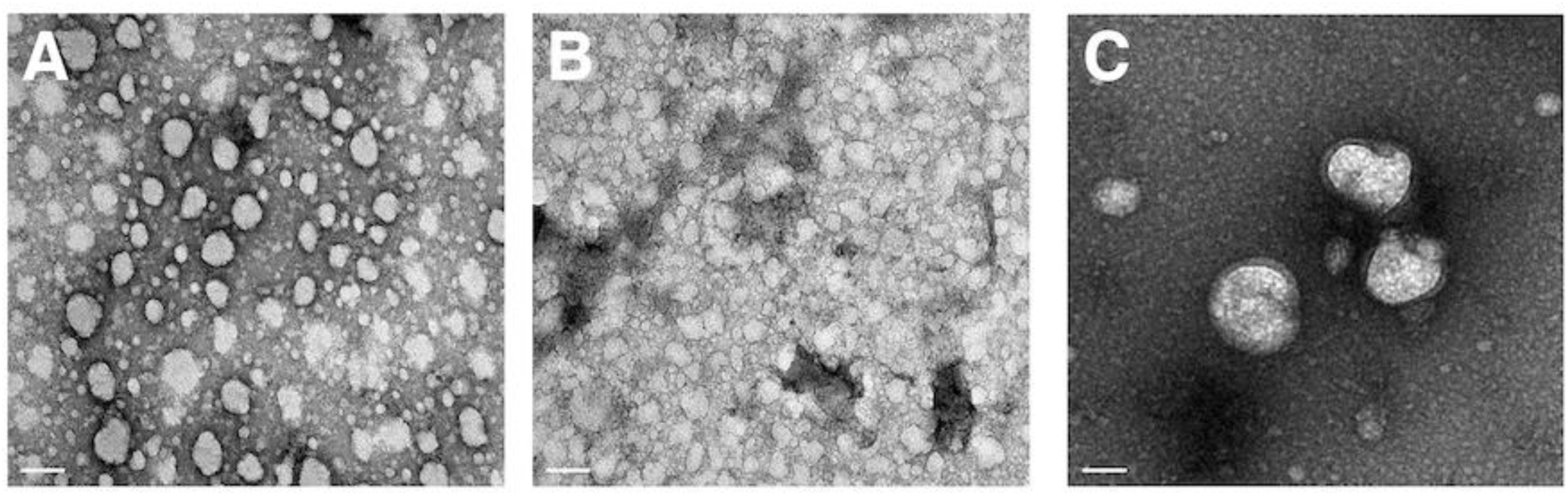
Transmission Electron Microscopy of Isolated Exosomes. Representative electron micrographs of plasma exosomes isolated by Total Exosome Isolation reagent of (A) non-pregnant and (B, C) GD14.5 exosomes. Scale bar represents 100nm.

**S2 Fig.**
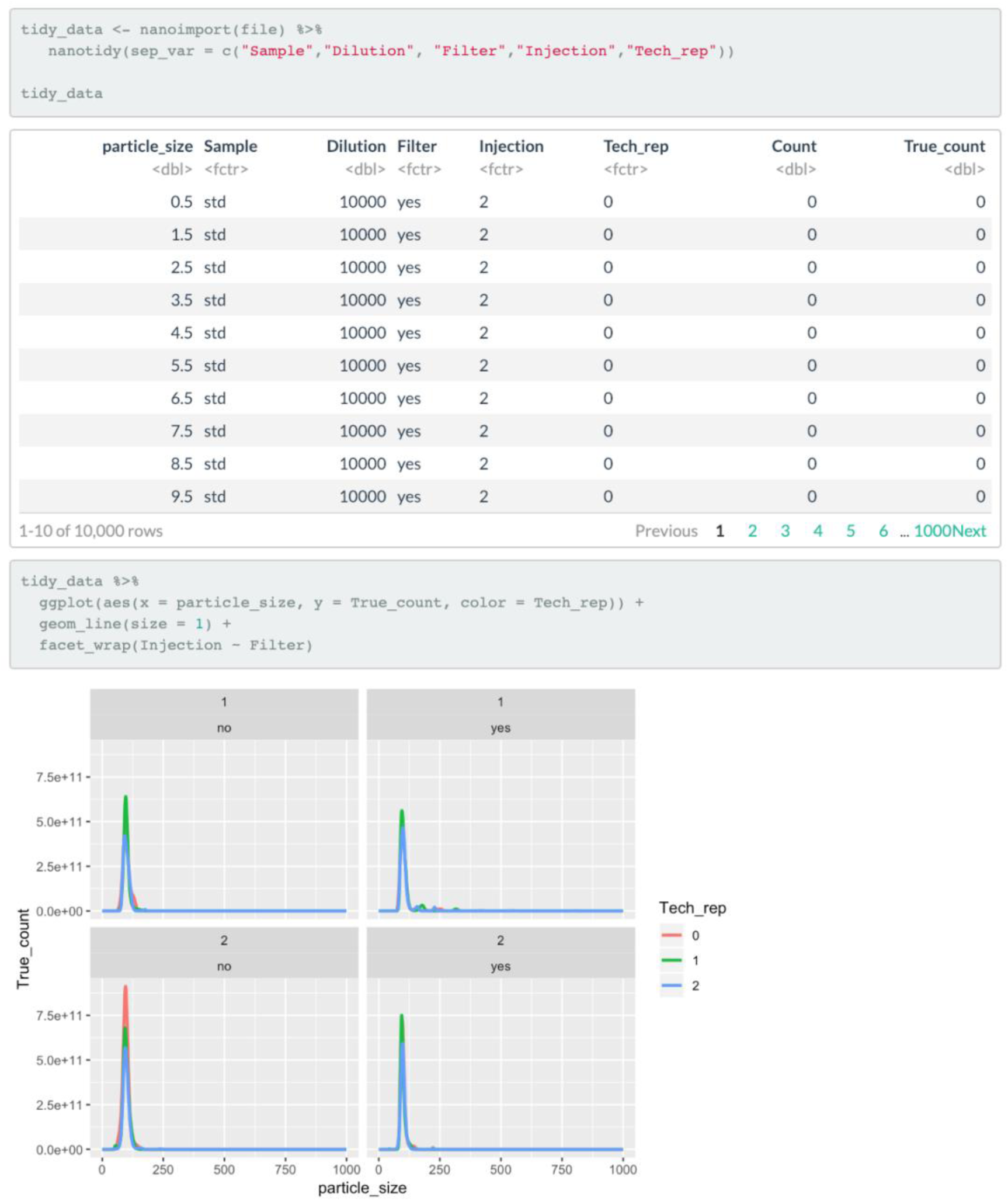
Polystyrene bead data analysis. Sample workflow of importing data into R using tidyNano functions.

**S3 Fig.**
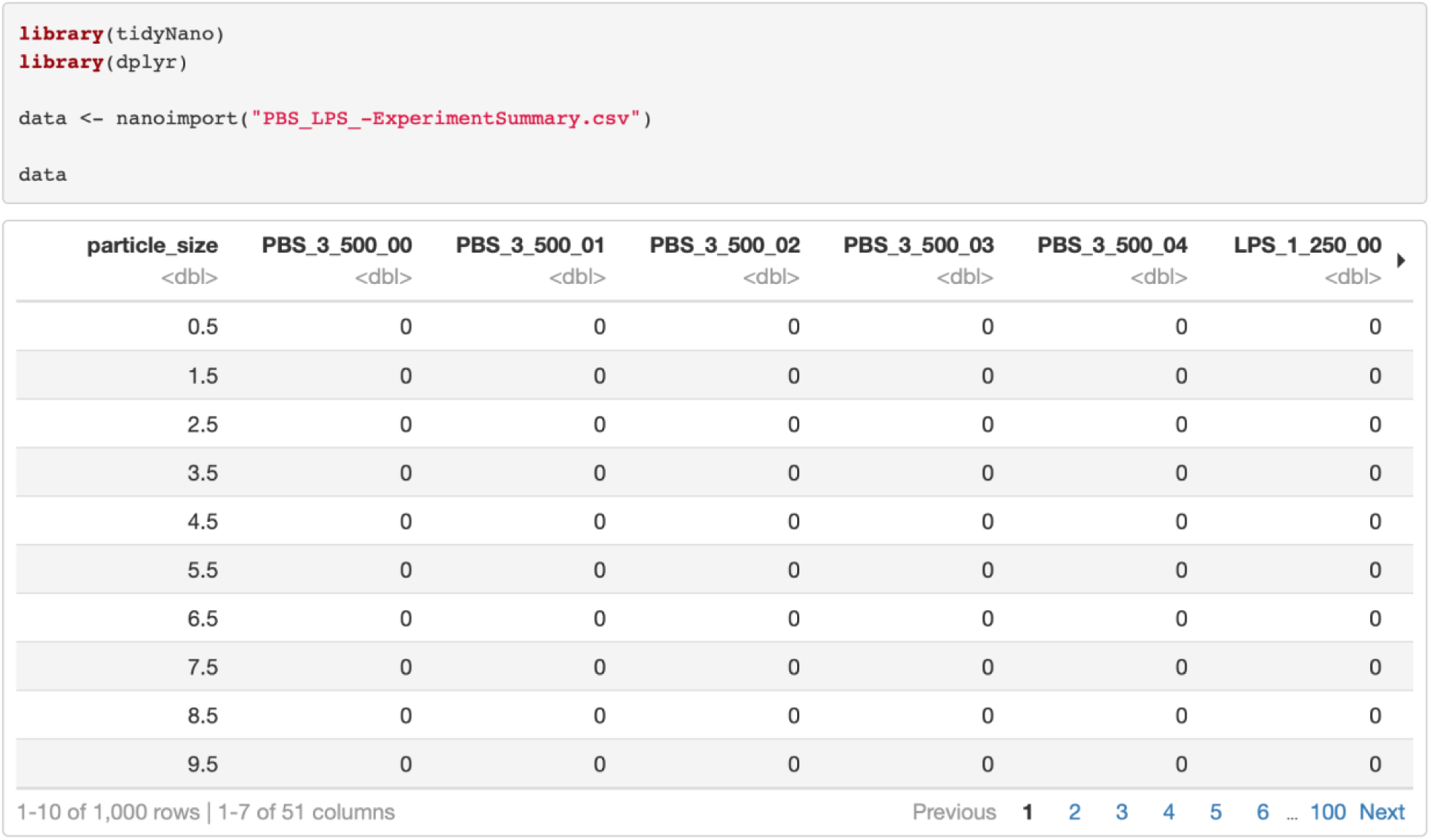
NTA data import with nanotidy(). Raw count data from NTA .csv files can be extracted and imported into the R environment with the nanotidy() function.

**S4 Fig.**
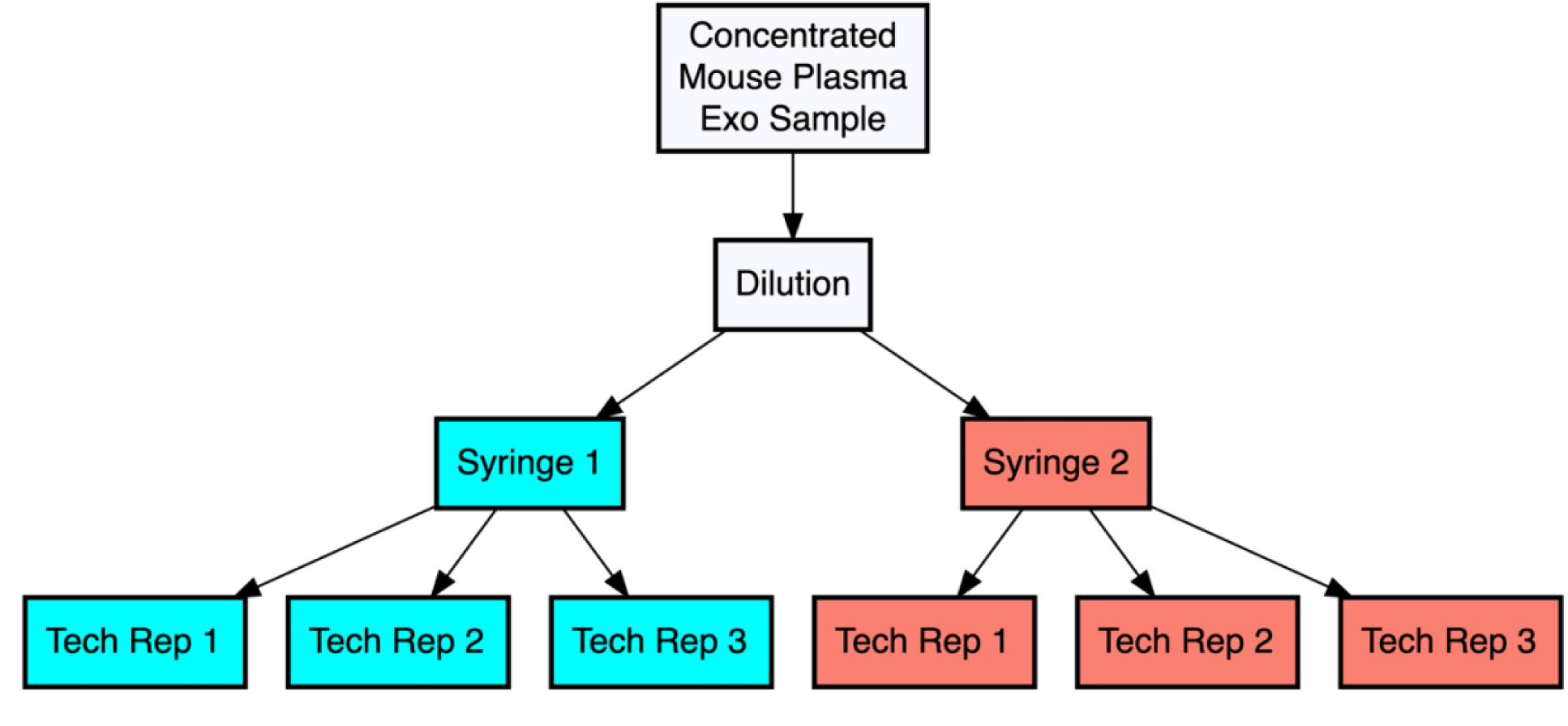
Schema of experimental design. Each plasma exosome sample was diluted, separated into two separate syringes, and was measured by NanoSight through recording of three 30-second videos.

**S5 Fig.**
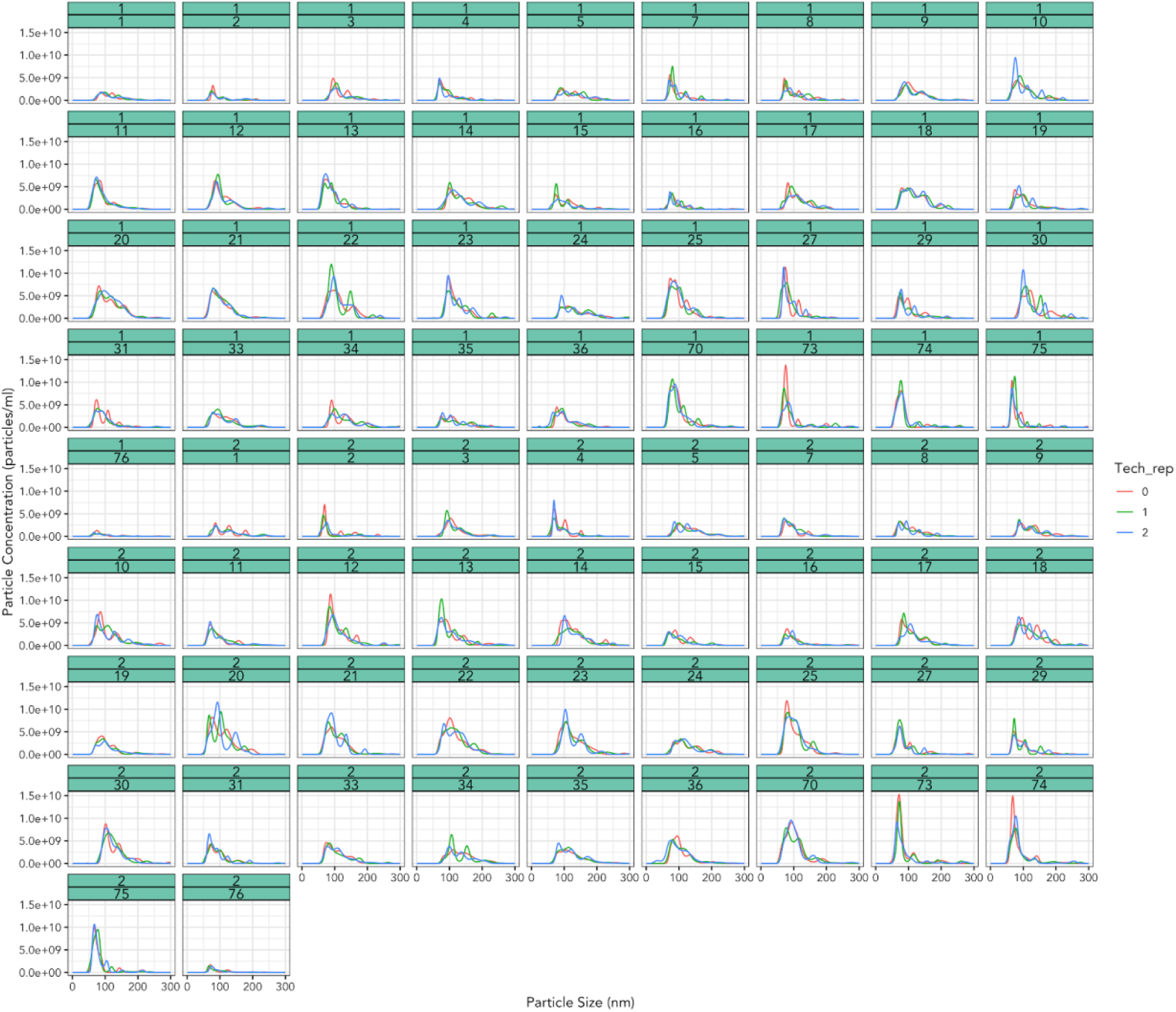
Visualization of samples with technical replicate data. Faceted plot of plasma exosome size and particle concentration of all samples (n=76) in experimental study where each sample was tested twice with three technical replicates.

**S6 Fig.**
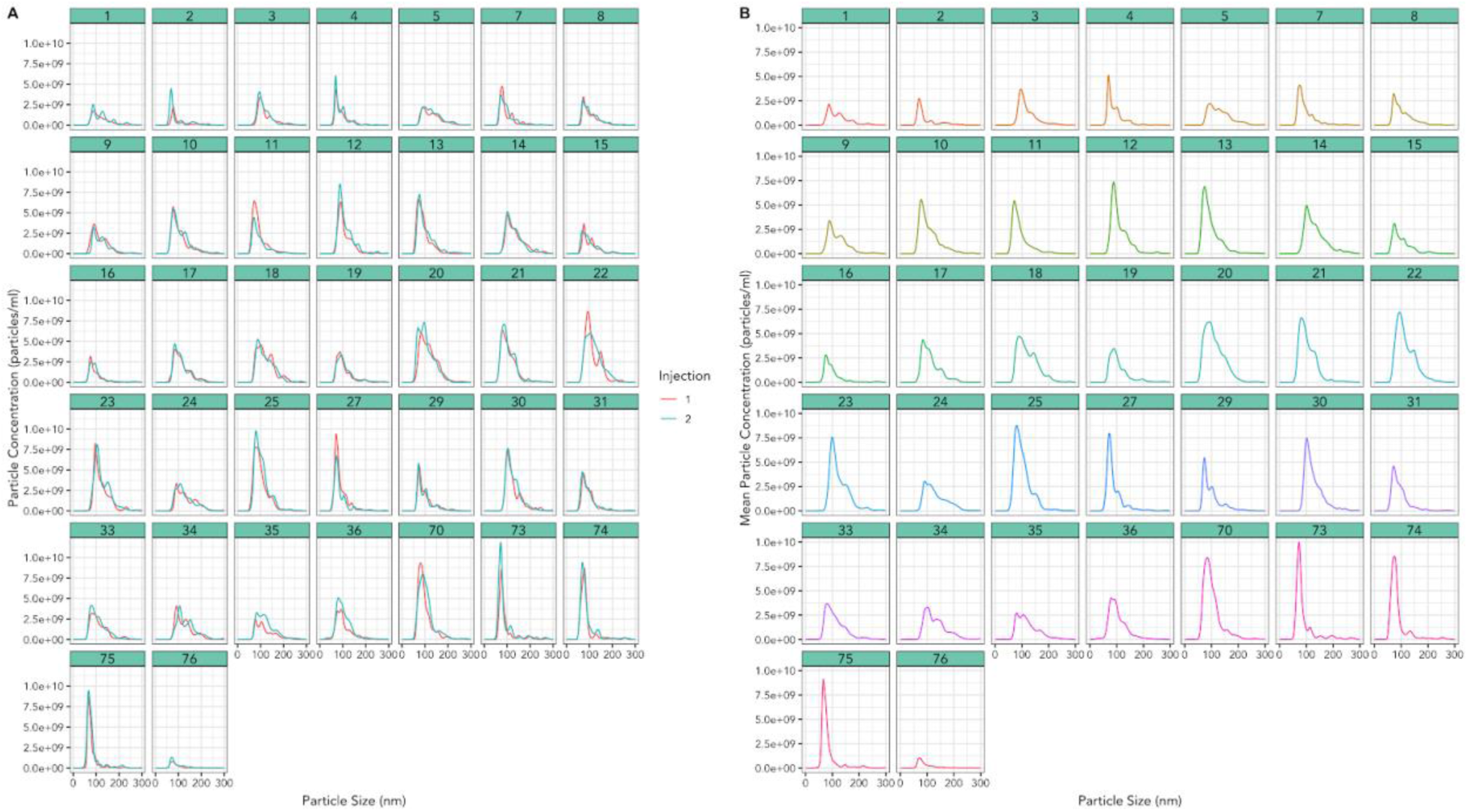
Visualization of samples with nanolyze(). Each summary output of the nanolyze() aggregation function can be visualized. (A) Plot of sample mean particle concentration from three technical replicate measurements (n = 76). (B) Plot of mean injection data from two syringe injection measurements (n = 76).

**S7 Fig.**
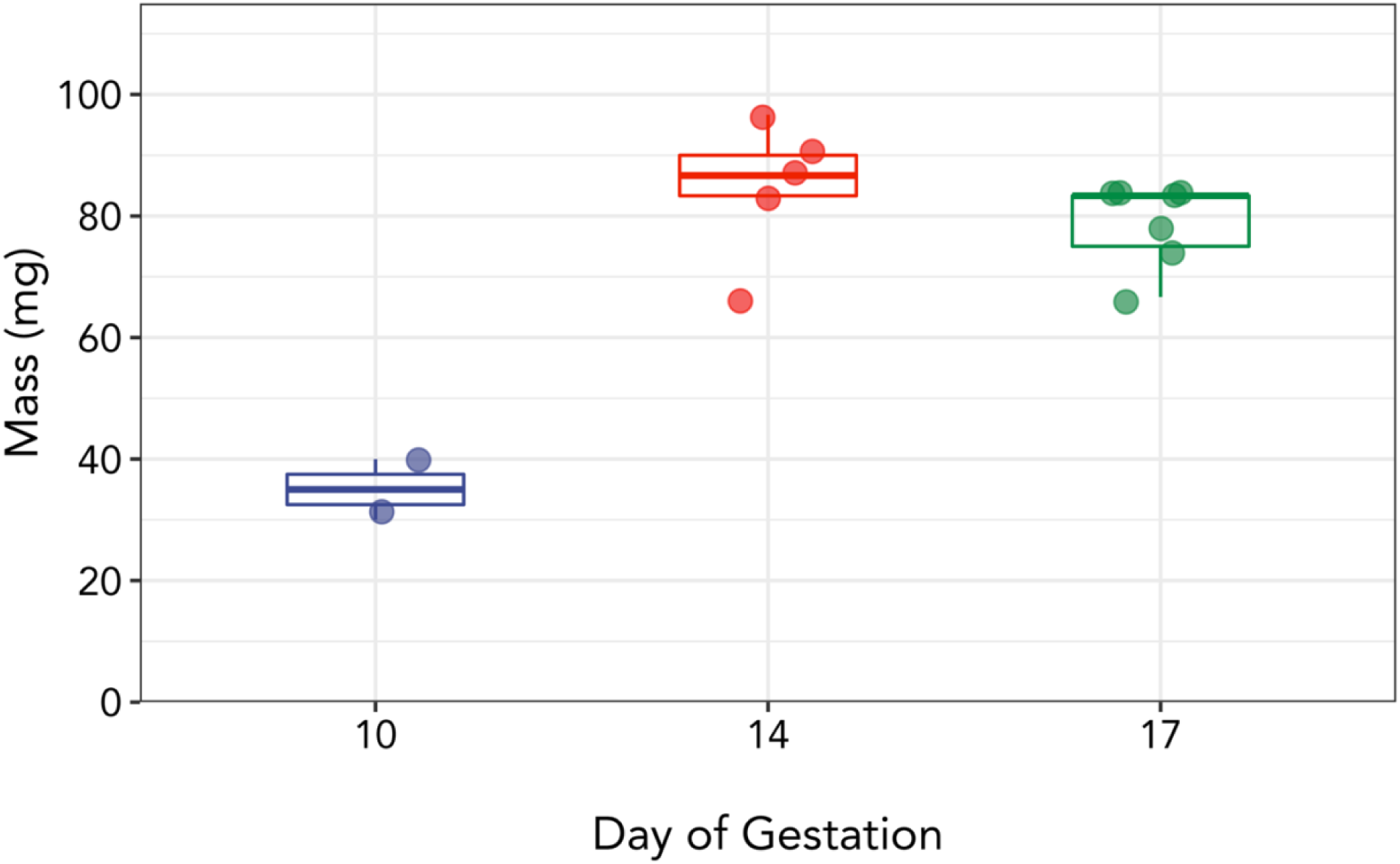
Placental mass across gestation. Placental mass across gestation in WT mated C57B/6 mice. Points represent individual placentas.

**S8 Fig.**
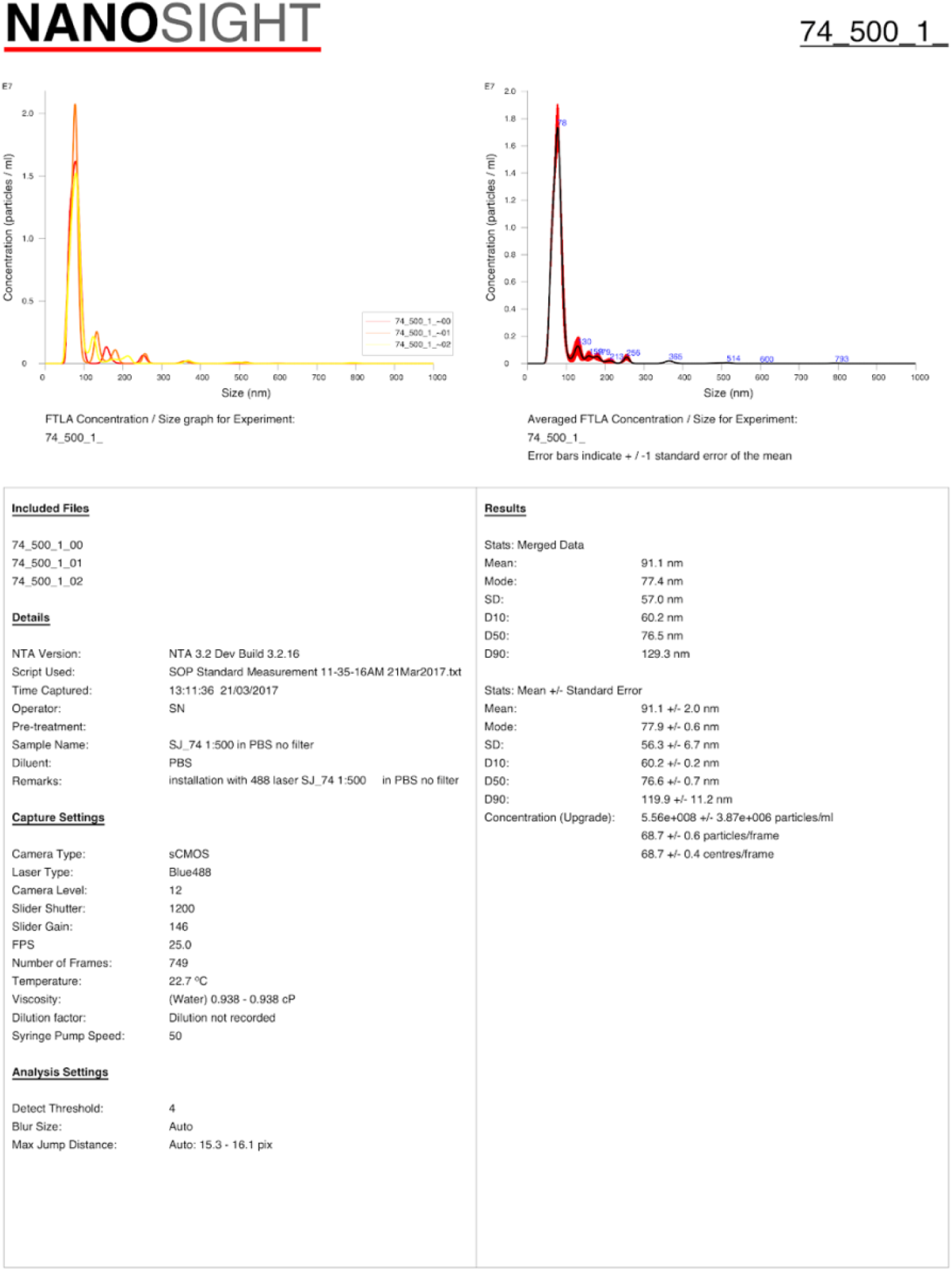
Sample NTA Summary PDF file.

